# *SAIBR*: A simple, platform-independent method for spectral autofluorescence correction

**DOI:** 10.1101/2022.01.19.476881

**Authors:** Nelio T.L. Rodrigues, Tom Bland, Joana Borrego-Pinto, KangBo Ng, Nisha Hirani, Ying Gu, Sherman Foo, Nathan W. Goehring

**Affiliations:** Francis Crick Institute, London, NW1 1AT, UK; Institute for the Physics of Living Systems, University College London, UK; Randall Centre for Cell and Molecular Biophysics, School of Basic and Medical Biosciences, King’s College London, London, SE1 1UL, UK

**Keywords:** autofluorescence correction, *C. elegans*, starfish, S. pombe, Fiji Plugin

## Abstract

Biological systems are increasingly viewed through the lens of mathematics, physics, and systems approaches that demand accurate quantification of gene expression and local protein concentrations. Such approaches have benefited greatly from the revolution in genetic engineering sparked by CRISPR/Cas9. By facilitating the tagging of genes at their genomic loci, CRISPR/Cas9 allows us to use fluorescence to monitor proteins that are expressed at or near endogenous levels under native regulatory control. However, due to their typically lower expression levels, quantitative experiments using endogenously-tagged genes can run into limits imposed by autofluorescence (AF). AF is often a particular challenge in the illumination bands occupied by the most efficient fluorescent proteins (GFP, mNeonGreen). Stimulated by our work in *C. elegans*, we describe and validate ***S**pectral **A**utofluorescence **I**mage correction **B**y **R**egression* (SAIBR), a simple, platform-independent protocol, and associated GUI-based FIJI plugin to correct for autofluorescence using standard filter sets and illumination conditions. Fully validated for use in *C. elegans* embryos and tested in diverse systems, including starfish oocytes and fission yeast, SAIBR achieves accurate quantitation of fluorophore signal and enables reliable detection and quantification of even weakly expressed proteins. Thus, SAIBR provides a highly accessible, low barrier way to incorporate AF correction as standard for researchers working on a broad variety of cell and developmental systems.

**Summary Statement:** Implemented as an easy-to-use Fiji Plugin, SAIBR provides effective autofluorescence correction for cells and tissues using standard imaging conditions.

## Introduction

Owing to its highly reproducible development and simple geometry, *C. elegans* has emerged as an ideal system for quantitative analysis of symmetry-breaking (Gross et al., 2019; Lang and Munro, 2017), cell division (Pintard and Bowerman, 2019), cell and tissue mechanics (Zhang et al., 2010), and cellular decision making (Barkoulas et al., 2013). Further, there is a wealth of endogenously-tagged genes of interest allowing live quantitative imaging of protein networks operating at native expression levels (Dickinson and Goldstein, 2016). However, despite being transparent, *C. elegans* exhibits significant intrinsic autofluorescence (AF) produced by a variety of cellular constituents and AF can be observed across all stages of worm development (Croce and Bottiroli, 2014; Pincus et al., 2016). It is most prominent when using blue and ultraviolet excitation wavelengths and thus poses problems when using standard GFP illumination conditions (Heppert et al., 2016). Consequently, there is a need for efficient and easily implemented methods for AF correction.

A number of general strategies have been sought to correct for AF. One approach is to experimentally suppress AF. One can use chemical compounds to specifically reduce or quench AF background, or even pre-bleach samples prior to fluorophore addition, though these methods tend to be restricted to fixed samples (Billinton and Knight, 2001; Cowen et al., 1985; Neumann and Gabel, 2002). In *C. elegans*, *glo* mutants exhibit reduced formation of autofluorescent gut granules, though because *glo* mutants affect normal worm physiology, they may complicate analysis (Hermann et al., 2005).

Another approach has been to optimize the combination of fluorophores, excitation light sources and emission filters to maximize the separation between fluorophore signal and AF (Billinton and Knight, 2001). These strategies usually require specialized imaging setups, and the narrow emission bands often required can limit the signal being captured. Such approaches may also restrict the choice of fluorophores. In *C. elegans*, the overlap of the excitation and emission spectra of AF with commonly used green fluorescence proteins, such as GFP or mNeonGreen (mNG), makes this approach difficult to achieve in practice. Nevertheless, some success has been achieved with specialized filter sets (An and Blackwell, 2003; Teuscher and Ewald, 2018) or yellow shifted excitation, which is compatible with mNG, but avoids the AF excitation peak (Heppert et al., 2016). One can also avoid the AF excitation peak by using red-shifted RFPs such as mCherry and mKate2. However, compared to GFP and mNG, red fluorophores are less optimal due to reduced quantum yield, lower brightness and enhanced photobleaching (Heppert et al., 2016; Shaner et al., 2004; Shcherbo et al., 2009).

Techniques such as fluorescence lifetime and spectral imaging can resolve overlapping fluorescent signals based on their distinct fluorescent lifetimes or spectral characteristics (Billinton and Knight, 2001; Kumar et al., 2009; Mansfield et al., 2005). AF typically exhibits fluorescence characteristics that are distinct from other fluorophores and, therefore, it can often be separated out much as one would an additional fluorophore. Such approaches have been useful in compensating for the high levels of gut autofluorescence in *C. elegans* (Shi and Grant, 2015). However, such techniques require specialized instruments and analytical tools, to which some may not have access or which may be incompatible with particular experimental workflows.

An alternative, but related approach to spectral unmixing is so-called ‘AF subtraction’ (Alberti et al., 1987; Billinton and Knight, 2001; Steinkamp and Stewart, 1986). First proposed for flow cytometry, this rather simple methodology takes advantage of the fact that AF typically exhibits much broader emission spectra than fluorophores such as GFP. One can therefore quantify AF in a given sample using an AF-reporting channel and subtract it from the signal measured in the fluorophore channel (Roederer, 2002). The advantages of this approach is that it is relatively straightforward to implement, it uses commonly available light sources and excitation/emission filters, and it does not require significant prior knowledge about fluorescence spectra beyond identifying a channel that is relatively specific for AF. Variations of this basic technique have since been applied in a variety of contexts, including cell-based monitoring of gene expression and AF correction of fluorescence in situ hybridization samples (Chen et al., 2019; Davis et al., 2010; Lichten et al., 2014; Roederer and Murphy, 1986; Szollosi et al., 1995).

Here we demonstrate that AF subtraction is a powerful method for autofluorescence correction during fluorescence imaging of *C. elegans* embryos. Notably, our implementation achieves results on par with more specialized imaging modalities, correcting both for bulk whole embryo fluorescence as well as for spatial AF variation. It enables reliable optimization of fluorescence signal from even very weakly expressed endogenous GFP fusions, and is compatible with dual labeled GFP/mCherry samples, bringing substantial improvements to fluorescence signal quantification. At the same time, due to its use of standard GFP/RFP filters sets and its implementation via an easy-to-use Fiji plugin, this protocol, which we term **S**pectral **A**utofluorescence **I**mage correction **B**y **R**egression (**SAIBR**), is readily combined with a variety of imaging platforms and procedures, allowing integration of AF correction as a standard part of imaging workflows. Although developed with *C. elegans* in mind, the principles behind SAIBR are general. Consistent with this, we further demonstrate that SAIBR is readily applicable to a variety of other experimental systems and thus will be a useful tool for AF correction for the cell and developmental biology community.

## Results

To quantify the potential impact of AF on the specific detection of GFP in *C. elegans* embryos, we began by comparing the magnitude of AF signal obtained from unlabeled embryos with the signal obtained from embryos expressing GFP fusion proteins when imaged with standard GFP illumination settings (ex^488^/em^535/50^, hereafter *GFP Channel*). For this purpose we selected *C. elegans* lines expressing GFP fusions to one of three polarity proteins, PAR-6 and PAR-3, which localize to an anterior plasma membrane domain, and LGL-1, which localizes to the posterior plasma membrane. All genes were tagged at the endogenous loci. In these cases, AF accounted for ~40-90% of the observed signal in the GFP emission band (Figure 1A). Moreover, when we imaged unlabeled embryos, the AF signal in the GFP Channel was intrinsically variable. Not only was there substantial spatial variation in AF (Figure 1B), but also a nearly 2-fold variation in the overall magnitude of AF signal between embryos (Figure 1A, B). Thus, simply subtracting out mean AF signal obtained from unlabeled embryos will fail to account for both sources of variation. In the case of LGL∷GFP such mean AF subtraction would clearly lead to apparent negative concentrations in some embryos. Thus, if we wish to accurately quantify the expression and local subcellular concentrations of proteins using GFP fusions, particularly for genes exhibiting low-to-moderate expression, one requires a method for locally correcting AF on a per embryo, per pixel basis.

**Figure 1.**
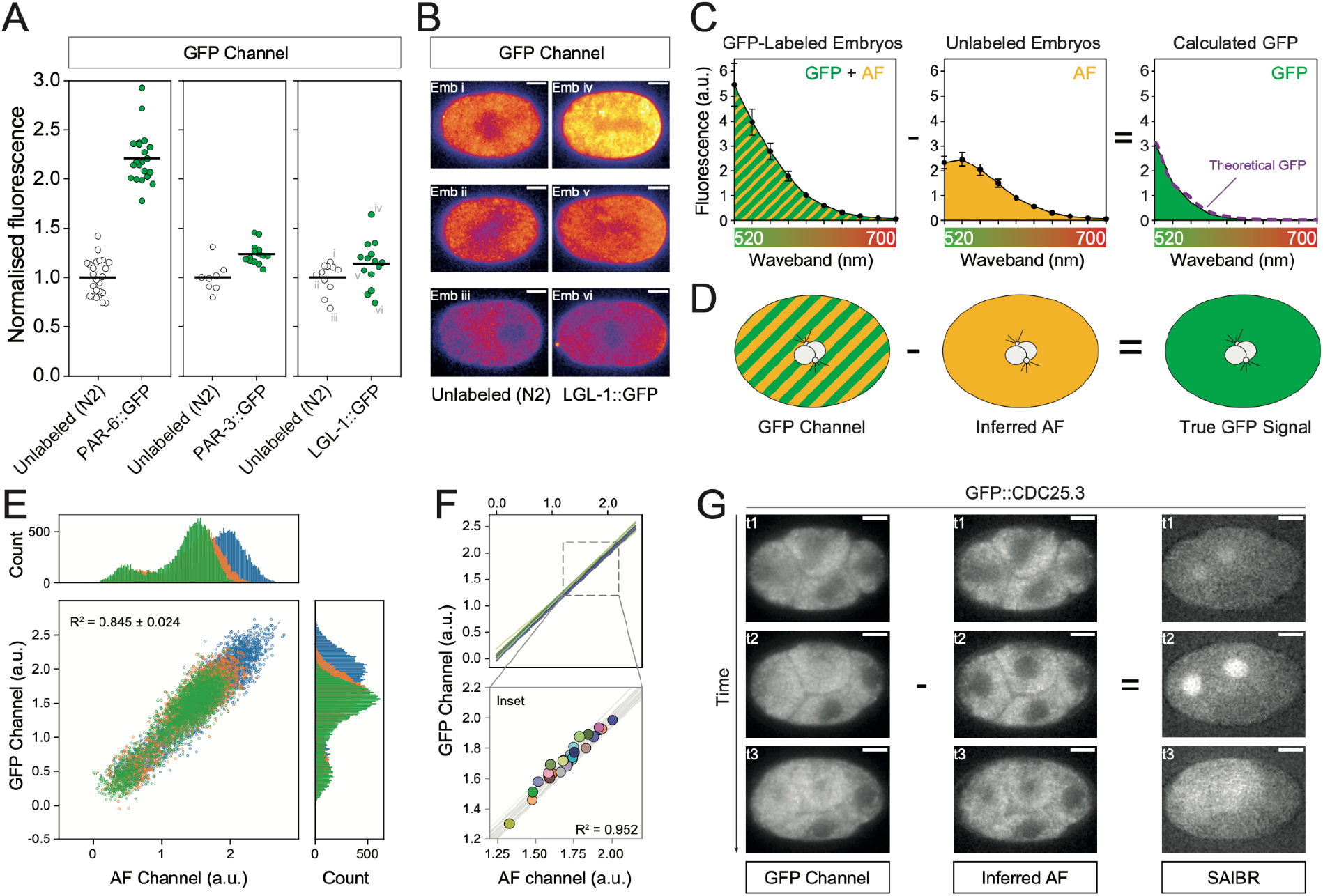
Autofluorescence in *C. elegans* is correlated across a broad spectrum of wavelengths. (A) Normalized fluorescence intensity for embryos expressing PAR-6∷GFP, PAR-3∷GFP, or LGL∷GFP (strains: KK1248, KK1216 and NWG0285, respectively) relative to unlabeled N2 embryo controls reveals that AF contributes substantially to total measured fluorescence signal. Data for individual embryos shown with mean indicated. (B) GFP Channel images of unlabeled N2 and LGL-1∷GFP embryos captured using identical parameters. Embryos taken from dataset in (A). (C) Subtraction of the fluorescence emission spectrum of unlabeled embryos (AF signal) from that of GFP∷PAR-6 expressing embryos yields a calculated GFP spectrum that is indistinguishable from the expected theoretical GFP spectrum (purple) demonstrating AF and GFP signals are additive. Fluorescence emission from embryos was measured in 20 nm wavebands with midpoints from 520 to 700 nm. Mean ± SD shown. N2, 5 embryos; PAR-6∷GFP, 5 embryos. (D) Schematic of SAIBR approach. Raw GFP Channel images consist of both GFP and AF signal. True GFP signal is obtained by subtracting inferred AF signal in the GFP Channel derived from AF measurement in a second channel. (E) Individual pixel values are well correlated between gaussian-filtered (radius = 1) AF and GFP Channels, and similar between embryos. Pixels from a region of interest comprising the entire embryo and a small section of surrounding background were used for the regression fit. A random selection comprising 10% all pixels shown color-coded per embryo. Histograms of intensity values for GFP and AF Channels shown for reference. (F) Comparison of per-pixel correlation with data obtained from whole embryo means. Lines indicate per-pixel regression from individual embryos as in (E). Inset shows overlay of mean whole embryo fluorescence values (circles). (G) Application of SAIBR to GFP∷CDC-25.3 expressing embryo. Subtraction of inferred AF signal reveals nuclear accumulation beginning at the start of the 4-cell stage and protein release at NEBD (t3). Scale bars = 10 μm.

### A simplified method for AF correction based on dual emission imaging

In principle, the emission signal for GFP and AF will be additive and thus if one has an independent measure of AF, one can subtract AF from the combined signal to obtain a value for GFP emission. Indeed, if we subtract the fluorescence emission spectrum of autofluorescence measured in unlabeled embryos from the spectrum obtained from embryos expressing PAR-6∷GFP, we almost perfectly recover the theoretical spectrum for GFP (Figure 1C). The challenge is therefore to find a method of quantifying AF directly in embryos also expressing GFP.

One strategy for quantifying AF takes advantage of the distinct spectral properties of AF that allow it to be quantified in an AF-reporting channel distinct from that used for measuring GFP, hereafter *AF Channel* (Alberti et al., 1987; Roederer and Murphy, 1986). One can use AF Channel measurements to correct for AF in the GFP Channel (Figure 1D).

In *C. elegans* embryos, AF peaks in the green-to-yellow wavelengths overlapping GFP, but extending further into the red (Figure 1C)(Heppert et al., 2016; Pincus et al., 2016). It can therefore be captured on a relatively selective basis through the use of a suitably red-shifted emission filter, such as those typically used for red fluorescent proteins. We therefore specify the AF Channel as ex^488^/em^630/75^. Note that in practice there will be a slight spillover of GFP signal into the AF channel, which could lead to overestimation of AF; however, because spillover is necessarily proportional to GFP amounts, it can be easily accounted for (see Methods). To establish an AF correction function between channels, we performed a linear regression on fluorescence pixel values obtained from images of unlabeled embryos captured in both the GFP and AF channels, where all signal is attributable to AF. We obtained strong linear correlations (R^2^ > 0.8) that were similar between embryos (Figure 1E). A nearly identical correlation was observed when we plotted the mean intensity values of entire individual embryos (R^2^ = 0.952, Figure 1F), indicating that the same correction function can account for both intra- and inter-embryo AF variation. Thus, we can use AF measurement in the AF Channel to accurately infer and therefore subtract out AF signal from the GFP Channel and obtain an accurate measure of ‘true GFP’ signal (Figure 1D).

As proof of principle, we captured images of embryos expressing a GFP fusion to CDC-25.3 from the endogenous locus. CDC-25.3 expression is repressed until early embryogenesis, reportedly becoming visible as embryos progress beyond the 8-cell stage (Tsukamoto et al., 2017). Using our AF correction method, we were able to observe clear nuclear localization already at the start of the 4-cell stage and accurately track its accumulation and release at NEBD at a time at which AF almost completely masked its expression in uncorrected images (Figure 1G).

We designated this protocol **S**pectral **A**utofluorescence **I**mage correction **B**y **R**egression (SAIBR). A schematic workflow is provided in Box 1, with additional details in Methods.

### AF correction using SAIBR

We next undertook a detailed quantitative analysis of the effectiveness of SAIBR in both unlabeled and GFP-labeled embryos. Applying SAIBR to unlabeled embryos effectively reduced observed embryo fluorescence in the GFP Channel to background, suggesting we accounted for nearly all AF signal in zygotes (Figure 2A). We then applied SAIBR to embryos expressing GFP fusions to LGL-1, PAR-3, or PAR-6 from the respective endogenous loci (Figure 2B-D). For both LGL-1∷GFP and PAR-3∷GFP, SAIBR revealed a clear peak in signal at the posterior and anterior plasma membranes, respectively, that was obscured by AF in uncorrected images (Figure 2B, 2C). Even when averaging cross-sectional membrane profiles across multiple embryos, membrane signal was difficult to discern in uncorrected data (Figure 2E, 2F, top). By contrast, SAIBR resolved membrane signal into a clear, well-defined peak (Figure 2E, 2F, bottom). Cytoplasmic signal also became substantially more uniform, which was most clearly visible in the suppression of a local fluorescence minimum in the embryo center due to AF exclusion by the pronuclei and mitotic spindle region. Improvements are also visible for PAR-6∷GFP-expressing embryos, though the magnitude of improvement is less striking due to the higher ratio of GFP to AF signal (Figure 2D, 2G). Note that similar results were achieved on both spinning disk confocal (Figure 2) and widefield microscopy (Supplemental Figure S1), confirming that the method is platform-independent.

**Figure 2.**
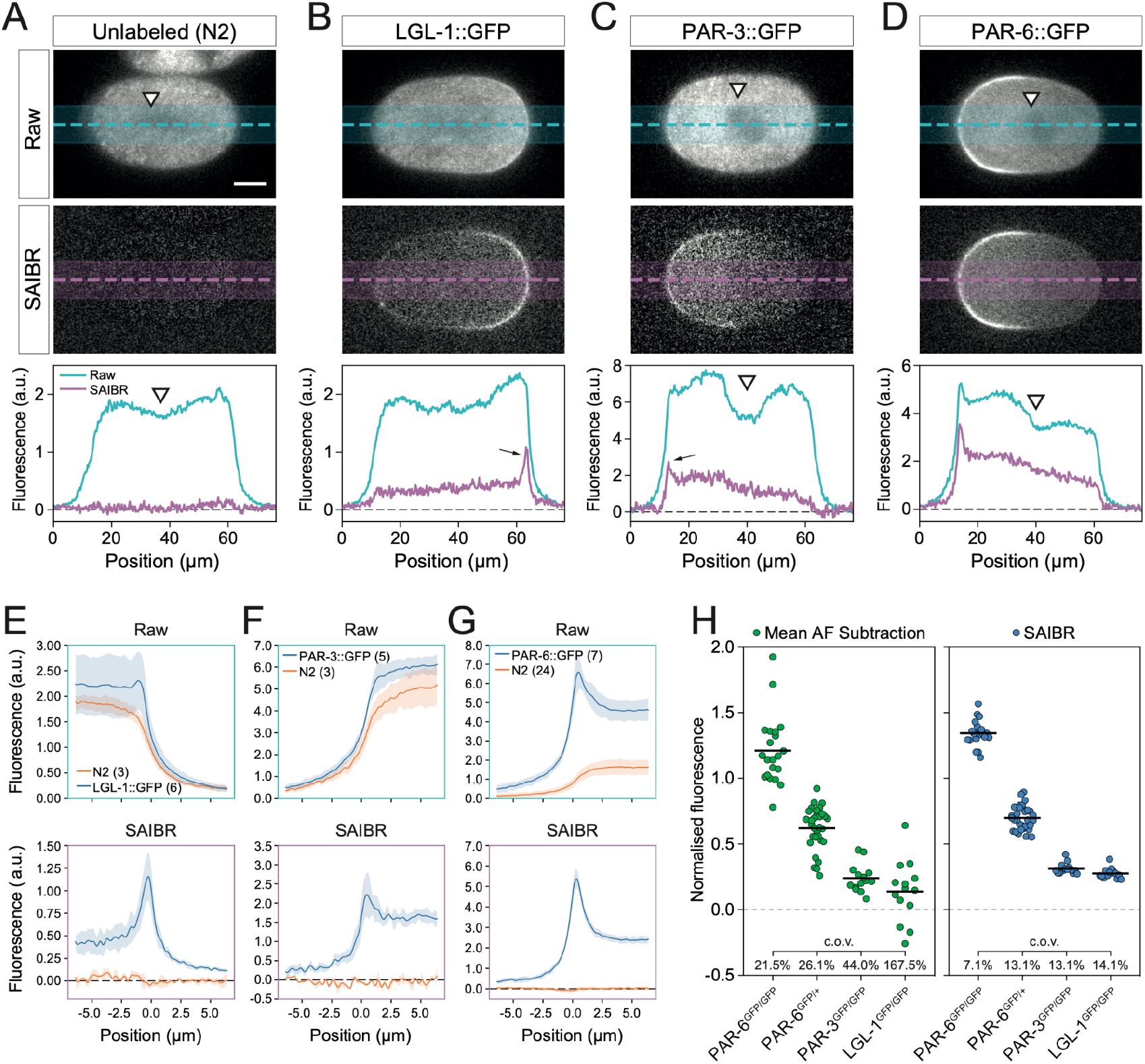
Quantitative analysis of GFP concentrations by SAIBR. **(A-D)** Raw (top) and SAIBR-corrected (middle) midplane images of zygotes expressing the indicated GFP fusions imaged in the GFP Channel as shown along with associated fluorescence linescan taken as indicated (bottom). Note SAIBR reduces GFP Channel signal to background in unlabeled embryos (A), and reveals prominent membrane localized signal in (arrows, B-C), which before correction is only obvious in PAR-6∷GFP embryos. Also note prominent depletion of signal in the pronuclear / spindle regions in GFP Channel images (white arrowhead) largely disappears in SAIBR images, suggesting it is due to local exclusion of AF signal. **(E-G)** Averaged membrane profiles taken from either raw (top) or SAIBR-corrected (bottom) images for LGL-1∷GFP (E), PAR-3∷GFP (F) and PAR-6∷GFP (G) shown relative to unlabeled N2 controls. Membrane position at x = 0μm. Mean ± SD shown. Number of embryos indicated (parentheses). Note membrane peaks are strongly enhanced for each GFP fusion expressing embryo, while N2 profiles are reduced to background in all datasets. **(H)** Quantitation of mean embryo GFP signal after correction by either mean AF subtraction (green) or SAIBR (blue) for embryos expressing PAR-6∷GFP in single (PAR-6^GFP/+^) or double copy (PAR-6^GFP/GFP^), PAR-3∷GFP (PAR-3^GFP/GFP^), or LGL-1∷GFP (LGL-1^GFP/GFP^). Mean fluorescence values for individual embryos shown with group mean and coefficient of variation (c.o.v.) indicated. Strains used: N2, KK1248, KK1216, NWG0285. Scale bars = 10 μm.

We next turned to quantification of total protein concentrations in embryos and compared SAIBR to a mean AF subtraction protocol (Mean AF Subtraction). For Mean AF Subtraction, we establish a mean AF signal in the GFP Channel based on fluorescence signal measured across multiple unlabeled embryos not expressing GFP and simply subtract this value from GFP Channel signal in GFP-expressing embryos. As a test, we used *C. elegans* lines expressing GFP∷PAR proteins described above as well as embryos that are heterozygous for the *par-6∷gfp* fusion and thus only half of the PAR-6 pool is labeled. For PAR-6∷GFP, both SAIBR and Mean AF Subtraction yield the expected 2:1 ratio of GFP signal between homozygous and heterozygous embryos (Figure 2H). However, by correcting for embryo-to-embryo AF variation, SAIBR substantially reduced the coefficient of variation (c.o.v.). We obtained similar results for PAR-3 and LGL-1 (Figure 2H). For LGL-1, in which the GFP signal was on the same order as variation in AF, the advantage of SAIBR was particularly striking. Whereas correction by Mean AF Subtraction resulted in negative values for GFP in embryos, the ability of SAIBR to suppress the effects of embryo-to-embryo variation in AF allowed it to achieve consistent and positive values for GFP signal in all embryos.

### Benchmarking against alternative strategies

To benchmark our method with other approaches, we used an alternative AF-minimization strategy in which we use a fluorophore compatible with wavelengths that minimize AF excitation. mNeonGreen (mNG) behaves similarly to GFP under standard GFP illumination settings (488 nm). However, due to a slight shift in its excitation spectrum, unlike GFP, it can be efficiently excited by a yellow-shifted laser line (514 nm) to substantially reduce AF (hereafter, *mNG Channel*, ex^514^/em^550/50^)(Heppert et al., 2016). Consistent with this observation, the magnitude of AF signal as measured in unlabeled embryos relative to total signal for embryos expressing mNG∷PAR-3 from the endogenous locus is reduced >2.5-fold in the mNG Channel relative to the GFP Channel (compare Figure 3A, 3B). We next compared the effectiveness of AF correction in three regimes, a standard regime using the GFP Channel and Mean AF Subtraction as described in the previous section, an mNG-specific regime using the mNG Channel and Mean AF Subtraction, and a regime using the GFP Channel but corrected by SAIBR. Plotting normalized corrected signal, we found that using either the mNG Channel or SAIBR regimes showed similar and substantial improvement in the variance of mean embryo fluorescence (Figure 3C), suppression of spatial varying cytoplasmic AF (Figure 3D), and accurate quantification of membrane signal (Figure 3E). Thus, SAIBR results in nearly identical improvement in signal quantitation and image quality as mNG-specific imaging conditions, but without the need for specialized laser lines or filter sets.

**Figure 3.**
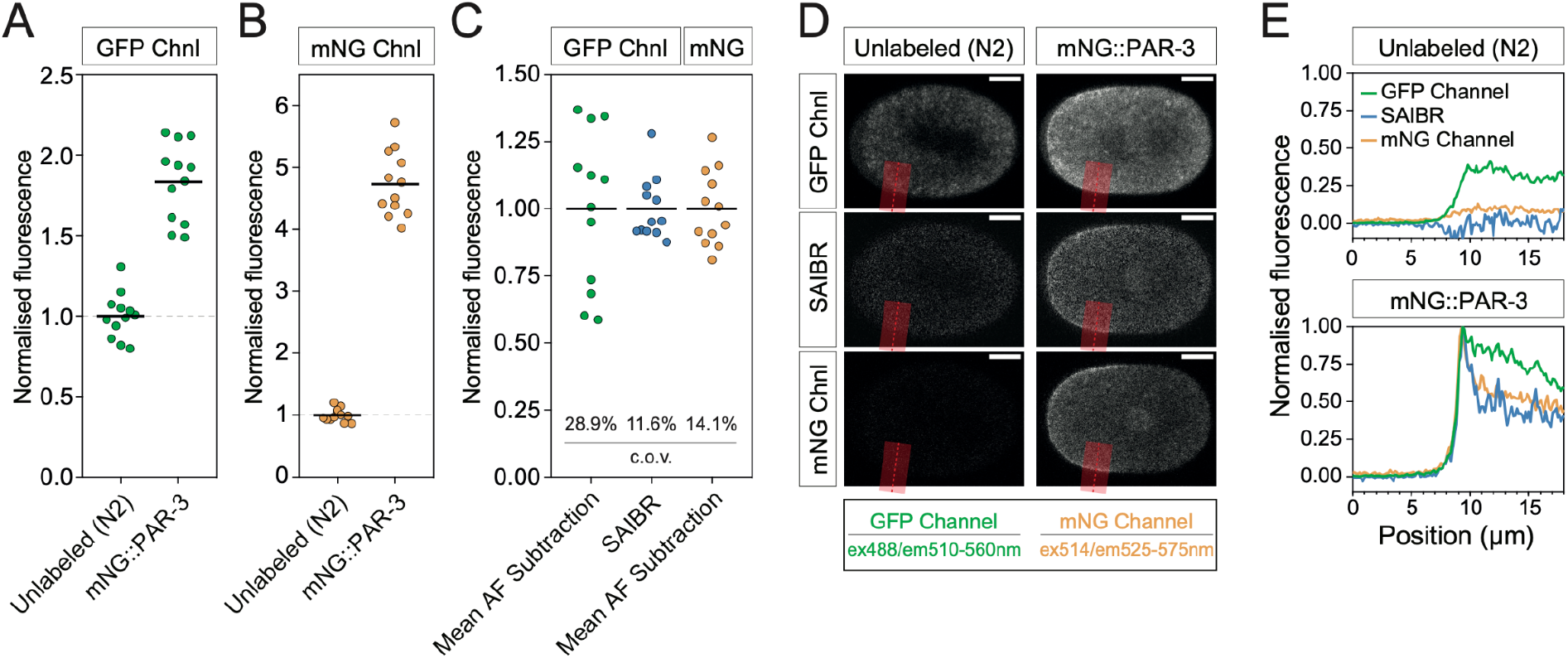
SAIBR achieves similar results to specialized excitation/emission imaging regimes. **(A)** Raw fluorescence signal using GFP illumination (GFP Channel, ex^488^/em^510-560^) for unlabeled (N2) or PAR-3∷mNG embryos (strain: NWG0189). Mean indicated. **(B)** As in (A) for mNG-specific illumination (mNG Channel, ex^514^/em^525-575^). **(C)** Comparison of mean normalized mNG fluorescence under three AF correction regimes as indicated. Note SAIBR and mNG-specific illumination yield similar reductions in CoV relative to standard GFP illumination with mean AF subtraction. **(D)** Midplane images of embryos excited imaged using the GFP Channel or mNG Channel and either left unprocessed (Raw) or subject to SAIBR as indicated. PAR-3∷mNG and unlabeled N2 shown for comparison. Scale Bars = 10 μm. **(E)** Plots of profiles taken perpendicular across the plasma membrane along the red line in embryos in (D) highlight similar performance of SAIBR and use of the mNG Channel in reducing AF.

### Extension of SAIBR to dual-labeled samples

Because SAIBR requires measurement of AF in a red-shifted emission channel, the use of dual fluorophore pairs such as GFP/mNG together with RFP/mCherry/mKate can introduce complications. Specifically, because red FPs (RFPs) are weakly excited by typical wavelengths used for GFP excitation, they will contribute to apparent AF, which is captured in the AF Channel. So long this contribution is low, i.e. for weak-to-moderate expression levels, it can be safely ignored (e.g. TH209, endogenous PAR-2∷mCherry, Figure 4A). However, at higher expression levels, the contribution of RFP signal to the AF Channel becomes significant (e.g. NWG0033, mCherry∷MEX-5, Figure 4A). At such levels, this bleedthrough signal induced a deviation in the mapping of AF between the AF and GFP Channels compared to N2 in proportion to the degree of RFP expression (Figure 4B) and leads to overcorrection for AF in the GFP Channel if not properly accounted for (e.g Figure 4F, left, 2-Channel SAIBR). In principle, this bleedthrough signal of RFP in the AF Channel is proportional to the concentration of RFP fusion and therefore is proportional to RFP signal captured using standard RFP illumination (*RFP Channel*, ex^561^/em^630/75^). We can therefore compensate for bleedthrough by using inputs from both the AF Channel and the RFP Channel to establish a revised 3-channel SAIBR correction function. To this end, we captured images of embryos expressing only RFP (no GFP) in the GFP, AF, and RFP Channels and performed a multiple linear regression in which AF in the GFP Channel is a function of signal in both the AF and RFP Channels (Figure 4C, 4D, Box 1). Applying 3-Channel SAIBR to embryos expressing PAR-6∷GFP with MEX-5∷mCherry eliminates oversubtraction and yielded measurements that were nearly identical to those for embryos expressing PAR-6∷GFP alone with no increase in data scatter, validating the expanded use of SAIBR to dual-labeled samples (Figure 4E, 4F). As a further test of its use in dual-labeled embryos, we used 3-Channel SAIBR to follow the relative localization of LGL-1∷GFP and PAR-6∷mCherry in time-lapse recordings of early embryos, which previously required multi-copy over-expression of LGL-1∷GFP (Beatty et al., 2010; Hoege et al., 2010)(Supplemental Movie 1).

**Figure 4.**
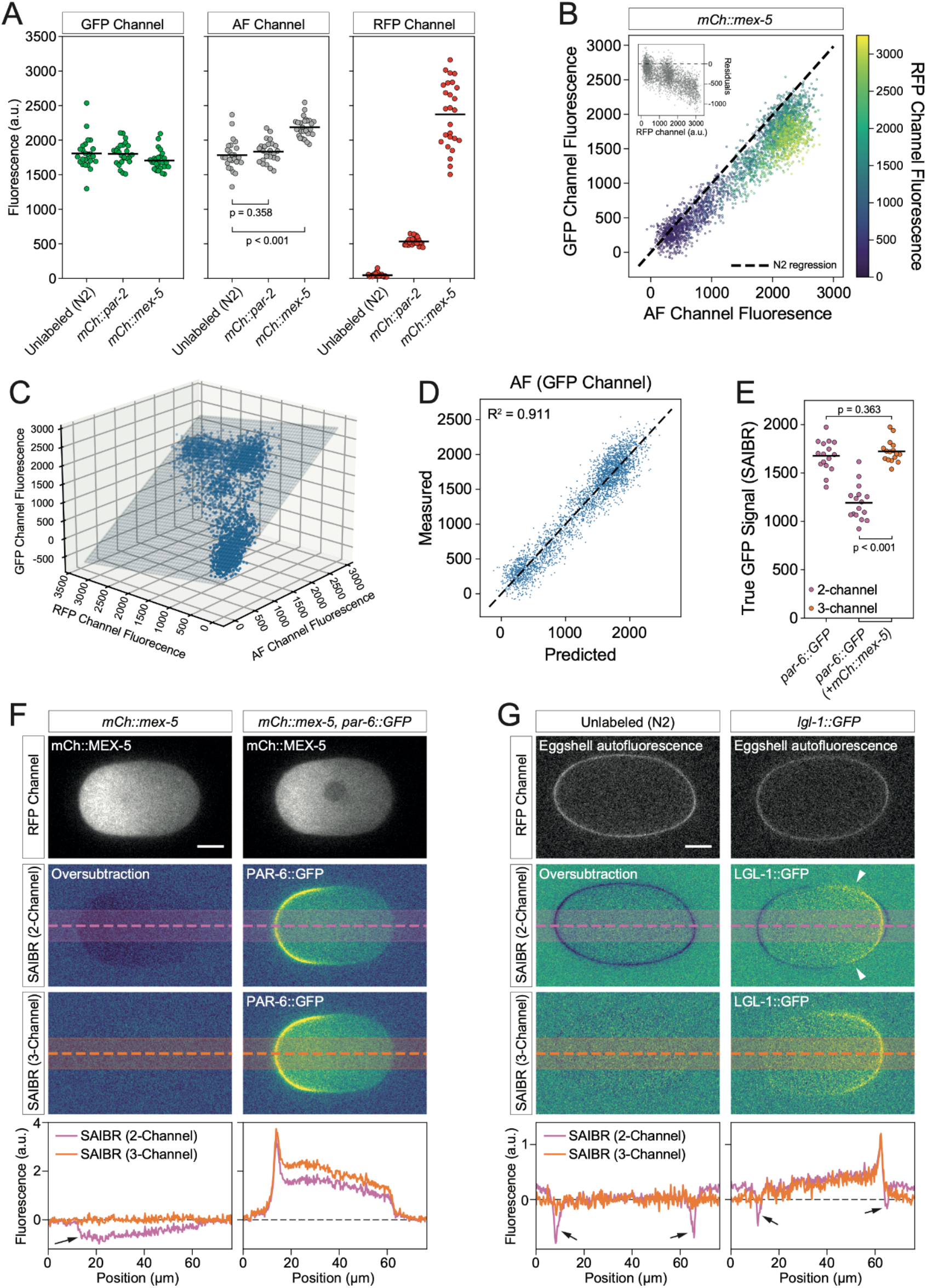
Spectral AF correction is compatible with dual color labeled samples. **(A)** Red fluorophore spillover increases signal in the AF Channel for highly expressed proteins. GFP, AF, and RFP Channel signal shown for TH209 (mCh∷PAR-2), or NWG0033 (mCh∷MEX-5) embryos relative to unlabeled N2. Note increase in the AF Channel is significant for NWG0033 due to higher levels of mCherry expression. p values determined using an unpaired t test. **(B)** mCherry spillover shifts the AF vs GFP correlation. Measured signal in gaussian-filtered (radius = 1) AF vs GFP Channel for a single NWG0033 (mCh∷MEX-5) embryo compared to wild-type N2 control, color-coded by RFP fluorescence (mCherry). Pixels from a region of interest comprising the entire embryo and a small section of surrounding background were specified and a random selection comprising 10% all pixels shown. Dashed line indicates AF vs GFP Channel correlation obtained from unlabeled N2 embryos. Inset shows residuals as a function of mCherry expression. **(C)** Multiple regression fit for 3-Channel SAIBR of AF signal in the GFP Channel as a combined function of signal in the AF and RFP Channels for an embryo expressing only mCh∷MEX-5. The same 10% sample of pixels is shown. **(D)** Predicted vs measured AF signal in the GFP channel based on the fit in (C). **(E)** 3-Channel SAIBR correction compensates for red fluorophore expression. Comparison of total PAR-6∷GFP signal in embryos co-expressing mCh∷MEX5 (strain: NWG0119) subject to either 2- or 3-Channel SAIBR compared to 2-Channel SAIBR applied to embryos expressing PAR-6∷GFP alone. Note overcorrection due to mCherry excitation by 488 nm illumination conditions results in underestimation of true GFP signal when using 2-Channel SAIBR. p values determined using an unpaired t test. **(F)** Images/profiles showing elimination of AF overcorrection in MEX-5∷mCherry expressing embryos. 2-Channel SAIBR using unlabeled N2 embryos for calibration results in oversubtraction that is visible as negative values (arrow) of NWG0033 (mCherry∷MEX-5) and in reduced cytoplasmic signal in NWG0119 (mCh∷MEX-5, PAR-6∷GFP). oversubtraction is eliminated with 3-Channel SAIBR using NWG0033 (mCh∷MEX-5) for calibration. **(G)** Simultaneous correction for embryo AF and egg-shell fluorescence. Eggshell fluorescence is strongly visible in the red fluorescence channel. Use of only 2-Channel SAIBR (calibration using only GFP and AF channels) results in oversubtraction of AF signal, visible as negative values in 2-Channel SAIBR images (arrows). This is eliminated by 3-Channel SAIBR that takes into account the RFP channel, improving signal of the LGL-1 posterior domain (strain: NWG0285). Arrowheads highlight regions of suppressed LGL-1 signal at the domain boundary due to eggshell-induced AF oversubtraction in 2-Channal SAIBR that does not occur in 3-Channel SAIBR..

Eggshell fluorescence is another issue in *C. elegans* embryos that sometimes arises and can potentially complicate SAIBR. Eggshell fluorescence is usually relatively minor in the GFP and AF Channels and can often be ignored. However, eggshell fluorescence is variable and may occasionally be significant for some methods of sample preparation and/or mounting (see Methods). Eggshell fluorescence has a distinct spectral profile compared to the AF we have discussed so far. Indeed, in many respects it behaves similarly to an RFP, though with a broader emission spectrum and hence a stronger bleed-through in the AF Channel. Pronounced eggshell fluorescence is particularly problematic when quantifying fluorophore signal at the plasma membrane because eggshell fluorescence will result in oversubtraction of AF in regions where the membrane and eggshell are in contact (Figure 4G). However, similar to an RFP, it could be compensated for by treating it as a red fluorophore and applying 3-Channel SAIBR (Figure 4G). It is important to note that because the emission spectrum of the eggshell is distinct from RFPs, one cannot simultaneously correct for both eggshell and RFP in the same samples.

### SAIBR in late C. elegans embryos and larvae

To expand the applicability of this method, we extended our analysis to late embryos (post-gastrulation) and larval stages. AF is known to be increasingly problematic in intestinal cells as *C. elegans* development proceeds and autofluorescent gut granules are formed (Laufer et al., 1980). Note that in unlabeled 1.5-fold stage embryos, the cells of the embryo posterior were visibly brighter due to AF signal in this region (Figure 5A). This signal was largely eliminated by SAIBR (Figure 5A). We next examined the localization of LGL-1∷GFP (Figure 5B, 5C). Some membrane staining of LGL is visible on the plasma membrane in uncorrected images, but was often obscured by AF. Thus, the locations of cells in many areas were only visible due to the reduced AF signal in the nucleus, particularly in the developing intestine (arrows). With SAIBR, the membrane localization of LGL-1∷GFP was much more clearly resolved and the cytoplasmic signal significantly more uniform. Notably, basolateral membrane localization of LGL-1 in intestinal cells could be clearly seen, juxtaposed to apical PAR-6∷mCherry signal (arrowheads) as observed previously in over-expression lines (Figure 5C)(Beatty et al., 2010; Sallee et al., 2021).

**Figure 5.**
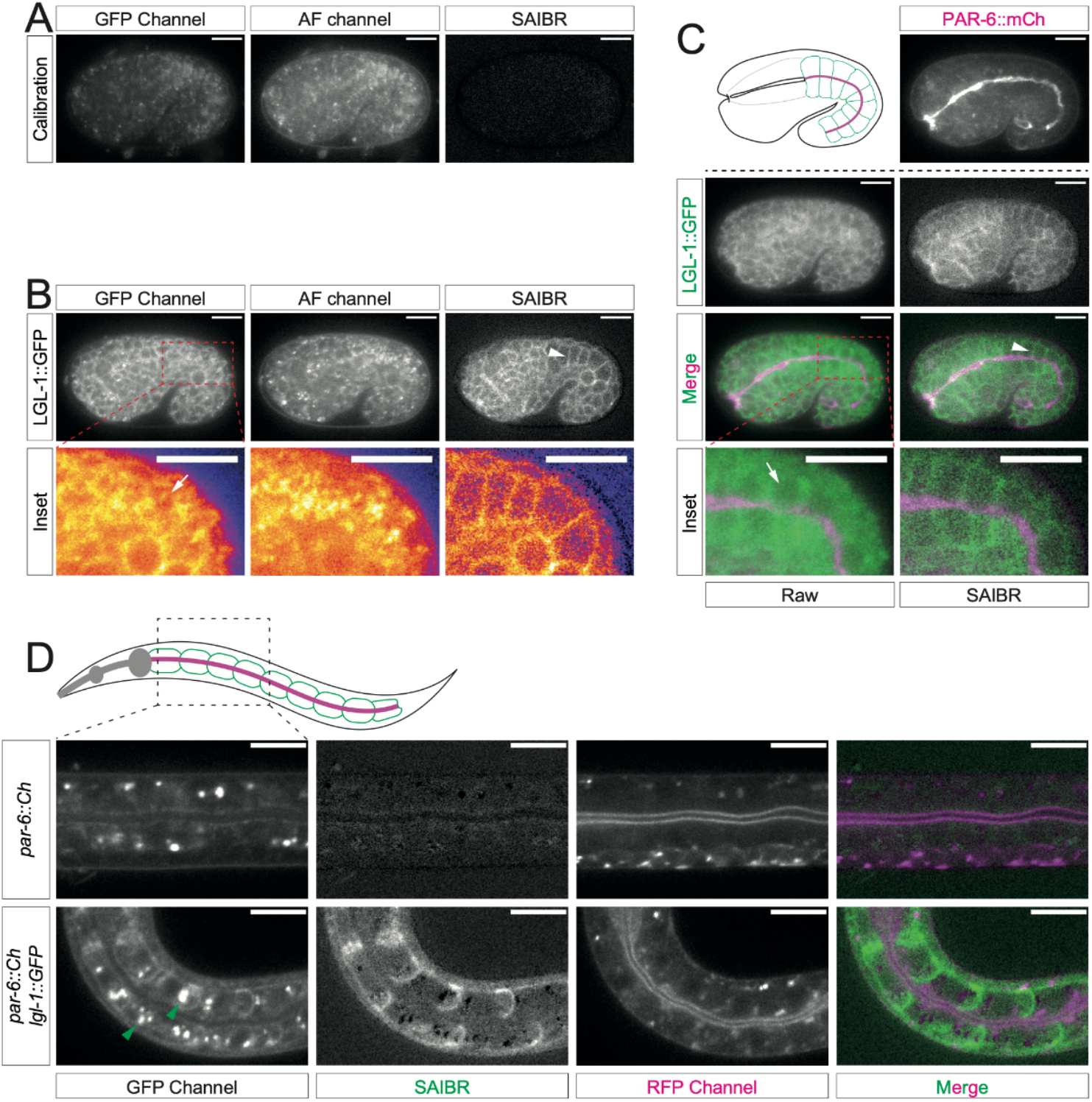
SAIBR suppresses autofluorescence background in late embryo and larval stages. **(A-C)** SAIBR reduces background in late stage *C. elegans* embryos. **(A)** Lateral view of a 1.5-fold embryo (Calibration, Strain BOX241) highlighting presence of substantial AF, particularly in the posterior near the nascent intestine, which is suppressed by SAIBR. **(B)** Posterior AF obscures membrane localization of LGL-1∷GFP, which is revealed by SAIBR. Below, Inset: zoom of posterior region (LUT - Fire). **(C)** SAIBR applied to a dual-labeled LGL-1∷GFP, PAR-6∷mCherry embryo reveals basolateral membrane localization of LGL-1. (top, left) Schematic of PAR-6 (magenta) and LGL-1 (green) localization in the developing intestine of a 1.5-fold stage embryo. (top, right) PAR-6∷mCherry labeling the apical domain of the intestine. (bottom) Paired uncorrected and SAIBR images shown alone, or merged with PAR-6∷mCherry visualized in the RFP Channel. Inset of merged images highlights posterior intestine. Note (B-C) show different planes of a single 1.5-fold stage embryo (Strain NWG0290). Arrows show nuclear void volume surrounded by fluorophore-specific and autofluorescence signals. Arrowheads show resolved LGL-1 membrane signal after image correction. **(D)** AF correction of *C. elegans* L1 larva reveals LGL-1∷GFP basolateral localization in the intestine. Still images of larval intestine expressing either PAR-6∷mCherry alone (Calibration, top row, BOX241) or LGL-1∷GFP and PAR-6∷mCherry (bottom row, NWG0290) shown for RFP (mCherry), GFP and SAIBR-corrected Channels along with a merge of RFP/SAIBR channels. Very bright puncta in the GFP Channel (green arrowheads) are gut granules which tend to be subject to overcorrection and therefore appear black in AF corrected images. Scale bars = 10 μm.

In L1 larva, gut granules are particularly prominent in intestinal cells in addition to the more diffuse AF signal characteristic of earlier embryos. Hence we were curious how SAIBR would perform (Figure 5D). When applying SAIBR, we found that a substantial fraction of granule fluorescence was over-subtracted, indicating that the fluorescence profile of mature gut granules differs from the more diffuse AF signal in the cytoplasm (Figure 5D, top). This emphasizes the problems associated with the presence of multiple, independently varying sources of AF. Nonetheless, despite modest oversubtraction of granules, SAIBR significantly improved visualization of LGL-1∷-GFP in these tissues (Figure 5D, bottom), revealing the expected basolateral pattern of localization in intestinal cells (Castiglioni et al., 2020). Thus, despite some oversubtraction of gut granule signal, our method can still improve visualization of weakly-expressed fluorophores in both late embryo and larval stages.

### A GUI-based FIJI SAIBR plugin enables simple AF correction in diverse systems

While the calibration and correction steps involved in SAIBR are relatively straightforward, we recognize that the need to implement such a workflow may limit widespread adoption. Therefore, to facilitate its use, we have implemented SAIBR as a simple GUI-based Fiji plugin which allows output of AF-corrected images in a few easy steps (summarized in Box 1). A detailed description of the plugin along with full instructions can be found together with sample datasets at https://github.com/goehringlab/saibr_fiji_plugin.

With the SAIBR plugin in hand, we solicited samples from a variety of experimental systems to validate its general suitability. To this end we obtained suitable sets of fluorescence images for two systems that exhibit autofluorescence. In starfish oocytes, bright autofluorescent cortical granules dominate the signal in the GFP channel, in this case in comparison to a relatively dim signal for the mother centriole (Figure 6A). The SAIBR plugin significantly suppressed the AF signal of granules, typically leaving the centriole clearly visible relative to the residual AF signal (Figure 6B). We observed a similar reduction in AF originating from vacuoles in the fission yeast, *S. pombe*, here shown relative to a mNG fusion to the ER/nuclear envelope-localized phosphatase component Nem1 (Figure 6C). These results confirm the potential broad applicability of SAIBR for AF compensation in cell and developmental systems.

**Figure 6.**
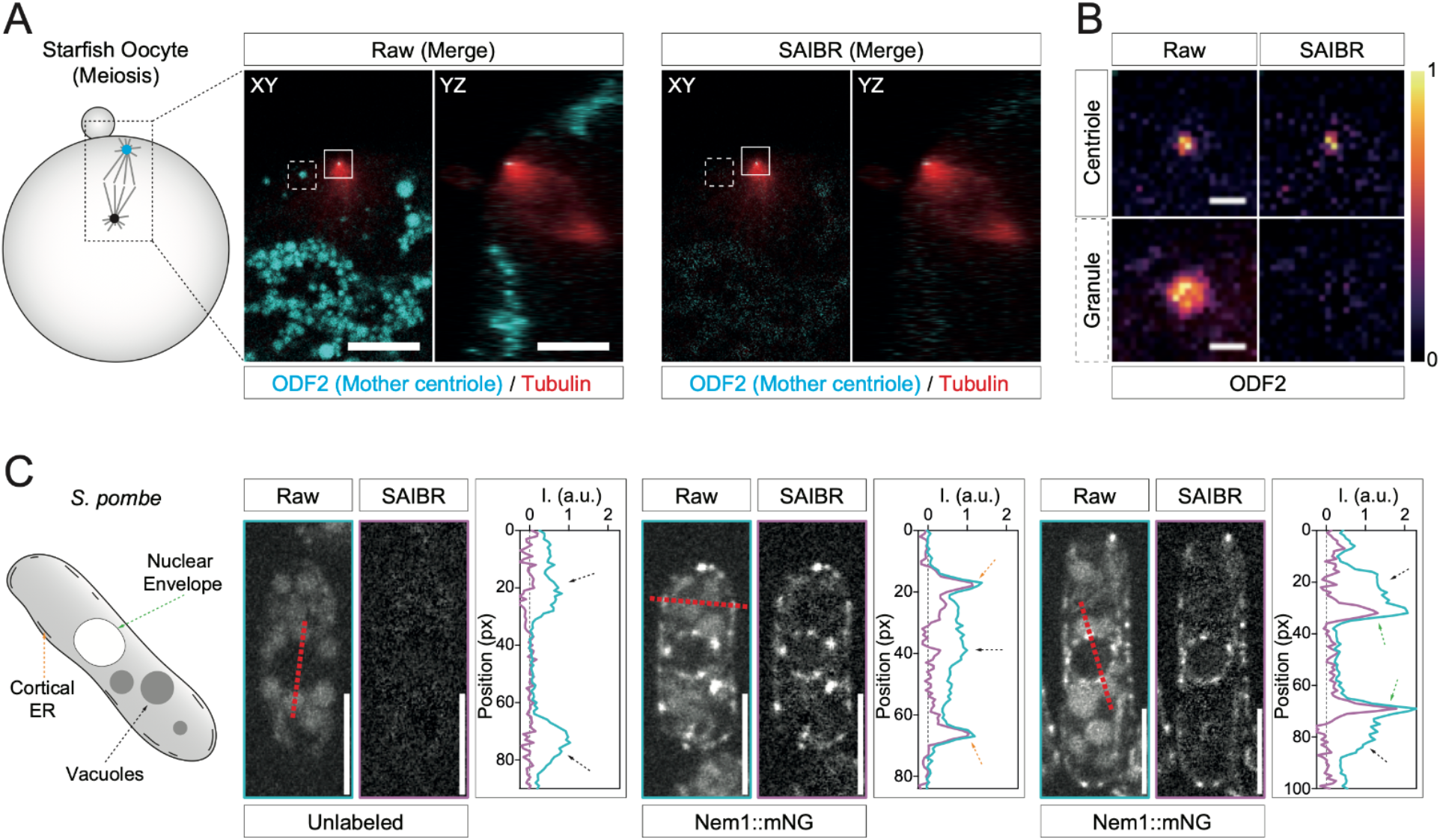
SAIBR effectively reduces AF in other model systems. **(A-B)** Suppression of cortical granule AF in starfish oocytes. **(A)** Image of a starfish oocyte labeled with ODF2∷GFP, rhodamine-tubulin is shown in XY and YZ planes during meiosis II. The spindle (tubulin, red) is perpendicular to the cell cortex with the mother centriole (ODF2, cyan) positioned at the spindle pole near the cell cortex. In raw, uncorrected images, yolk granules (cyan) dominate the signal in the GFP Channel (left, Raw). Yok granule signal is strongly suppressed by SAIBR leaving the ODF2-marked centriole as the strongest signal (right, SAIBR). Scale bars = 10 μm. **(B)** Scaled, single channel insets (10x zoom) of the ODF2-marked mother centriole and neighboring yolk granule from (A) highlighting specific suppression of yolk granule fluorescence relative to ODF2 (LUT - inferno). Scale bars = 1 μm. **(C)** Suppression of vacuolar AF in fission yeast cells. Raw (blue) and SAIBR-corrected (purple) midplane images of yeast cells from an unlabeled (left cell) or Nem1∷mNG-expressing strain (middle and right cells) imaged in the GFP channel. Quantitation of fluorescence linescans taken across cells as indicated (see red dashed lines). Note SAIBR reduces non-specific AF signal (mostly emitted from vacuoles - see black arrows) to background in the unlabeled cell, and reveals a prominent, well resolved Nem1-specific signal at the cortical ER (see orange arrows - middle cell) and nuclear envelope (see green arrows - right cell). Scale bars = 5 μm.

## Discussion

Here we describe and validate a simple and easy-to-use implementation for autofluorescence correction, SAIBR, provided as a FIJI plugin. A simplified form of spectral imaging, we leverage the typically broad AF spectrum to allow accurate estimation and correction of AF signal in the GFP Channel. SAIBR is platform-independent and relies only on commonly available GFP/RFP illumination sources and filter sets. Yet SAIBR yields similar performance to more specialized methods in both single and dual fluorophore labeled samples, enabling the visualization and accurate quantification of even weakly expressed proteins that are at the limits of detection above AF.

Our aim was to provide a tool that would enable regular and widespread use of AF correction by the research community as part of day-to-day investigation. Autofluorescence is a particularly common problem when imaging GFP in a number of systems with sources including yolk and cortical granules, extracellular matrix, and lysosomal compartments. While more complex solutions exist, as we have shown, SAIBR provides a simple and straightforward solution that is likely to work efficiently in many contexts. The ease of use of the SAIBR plugin and the generic imaging conditions required mean that it costs users little to test on their system of choice, allowing one to quickly determine whether a more complex approach is required.

The simplicity and platform independence of SAIBR allow it to be integrated into a variety of experimental workflows with minimal extra investment of time and resources. We envision that SAIBR and its potential to drive widespread adoption of routine AF correction should enable new experimental approaches. In the *C. elegans* embryo, for example, by allowing accurate AF correction on a per embryo basis, not only will this method provide improved measurement of protein concentration and subcellular distribution within cells, but it will also allow us to address questions related to protein dosage, such as assessing variability of protein expression and its potential effect on developmental pathways.

In principle, there is no need to restrict oneself to the wavelengths used here, which were chosen to solve the particular problem of GFP Channel autofluorescence. The technique itself only requires the identification of an appropriate channel that can be used to infer AF in the desired fluorophore-reporting channel and hence should be well separated from the channel used for fluorophore imaging.

At the same time, simplicity and flexibility come with certain tradeoffs. First, SAIBR requires identification of an AF channel that is reasonably well-isolated from the reporter channel for the fluorophore of interest. Second, it relies on the existence of an AF signal that exhibits a consistent correlation between AF and fluorophore channels and hence is not suited to samples with multiple, independently varying sources of AF. Third, SAIBR combines pixel noise from multiple channels, effectively amplifying salt-and-pepper noise in images, though one could in principle apply computational denoising strategies to reduce this effect (Krull et al., 2019). Finally, as implemented here, SAIBR requires one to capture images in at least two emission channels, effectively doubling sample illumination and minimum time intervals The time lag between frames may also lead to pixel mismatches between GFP and AF channels for samples exhibiting rapid motion. However, this last limitation can be bypassed with suitable optics to allow for simultaneous capture of multiple emission bands.

In summary, by combining ease of implementation and accurate AF correction with relatively few tradeoffs, we hope SAIBR will help facilitate widespread adoption of autofluorescence correction and enable more accurate quantification of the concentration and distribution of fluorescently-tagged proteins in live cells and tissues.

## Methods

### C. elegans - strains and maintenance

*C. elegans* strains were maintained on OP50 bacterial lawns seeded on nematode growth media (NGM) at 20°C under standard laboratory conditions (Stiernagle, 2006). Strains are listed in Supplemental Table S1. OP50 bacteria were obtained from CGC.

### C. elegans - strain construction

Mutation by CRISPR-Cas9 was performed based on the protocol published by (Dokshin et al., 2018). Briefly, tracrRNA (IDT DNA, 0.5μL at 100μM) and crRNA(s) for the target (IDT DNA, 2.7μL at 100μM) with duplex buffer (IDT DNA, 2.8μL) were annealed together (5 min, 95°C) and then stored at room temperature until required. PCR products containing the insert DNA sequence (GFP in this instance) and insert with an additional 130 bp homology to the insertion site were generated, column-purified (Qiagen, QIAquick PCR purification kit), mixed in equimolar amounts, denatured by heating to 95°C, and followed by annealing thorough slow cooling to room temperature to generate a pool of products with long single stranded DNA overhangs to act as the repair template. An injection mix containing Cas9 (IDT DNA, 0.5μL at 10mg/mL), annealed crRNA, tracrRNA, and the repair template was incubated at 37°C for 15 min and centrifuged to remove debris (15 min, 14,500rpm). Young gravid N2 adults were injected along with a *dpy-10* co-CRISPR injection marker (Arribere et al., 2014) and mutants identified by PCR and sequence verified. Resulting lines were backcrossed with N2s twice before use. Sequences available in Table S1.

### C. elegans - dissection and mounting for microscopy

For most experiments, early embryos were dissected from gravid hermaphrodites in 5-6 μL of M9 buffer (22 mM KH_2_PO_4_, 42 mM NaHPO_4_, 86 mM NaCl and 1 mM MgSO_4_) on a coverslip and mounted under 2% M9 agarose pads (Zipperlen et al., 2001). In some instances (Figure 1B, 1G, 2A, 2B, 2E, 3, and Supplemental Figure S1), to minimize eggshell autofluorescence that may be prominent with agarose mounts, embryos were dissected in 8-10 μL of egg buffer (118 mM NaCl, 48 mM KCl, 2 mM CaCl_2_ 2 mM MgCl2, 25 mM HEPES, pH 7.3), and mounted with 20 μm polystyrene beads (Polysciences, Inc.) between a slide and coverslip as in (Rodriguez et al., 2017).

To harvest late embryos (Figure 5A-C), gravid worms were allowed to lay embryos for 4-5h at 20°C. Embryos were collected and mounted in 8-10 μL of egg buffer supplemented with 18.8μm polystyrene beads (Polysciences, Inc.).

L1 larva (Figure 5D) were collected from plates where gravid adult worms were allowed to lay eggs for 12-13h at 20°C. Whole larva were then mounted between a 2% M9 agarose pad and coverslip in M9 containing 0.1μm polystyrene beads (Polysciences, Inc.) and 10 mM Levamisole to induce worm paralysis (Reich et al., 2019).

### C. elegans - Fluorescence microscopy

Unless specified otherwise, midsection confocal images were captured on a Nikon TiE with a 60x/1.40 NA oil objective, further equipped with a custom X-Light V1 spinning disk system (CrestOptics, Rome, Italy) with 50 μm slits, Obis 488/561 fiber-coupled diode lasers (Coherent, Santa Clara, CA) and an Evolve Delta EMCCD camera (Photometrics, Tuscon, AZ). Imaging systems were run using Metamorph (Molecular Devices, San Jose, CA) and configured by Cairn Research (Kent, UK). Filter sets were from Chroma (Bellows Falls, VT): ZT488/561rpc, ZET405/488/561/640X, ET535/50m, ET630/75m. For late embryo and larval imaging, a 1.5x magnification was applied (using TiE intermediate magnification switching).

Midsection wide-field fluorescence images were captured on a Nikon TiE with a 60x/1.40 NA oil objective, further equipped with a Spectra-X Light Engine (Lumencor, Inc., Beaverton, OR). Imaging systems were run using Metamorph (Molecular Devices, San Jose, CA) and configured by Cairn Research (Kent, UK). Filter sets were from Chroma (Bellows Falls, VT): ET490/20x, ET525/50m, ET632/60m.

To obtain the emission spectra shown in Figure 1C, embryos were imaged under 488 nm excitation at consecutive 20-nm wavebands over the range of 510-710 nm (yielding lambda stacks of 10 images per embryo). Wavelength-scans were performed on a Leica TCS SP8 inverted microscope (Leica Microsystems Ltd, Wetzlar, Germany), equipped with an Apo CS2 63x/1.40 NA oil objective and a HyD detection system. Imaging was managed with LAS X software (Leica Microsystems Ltd, Wetzlar, Germany), and acquisition was set at a scanning speed of 700 Hz with pinhole aperture set to 2 AU.

For midsection confocal imaging of mNG-expressing embryos and comparison between 488 nm and 514 nm excitation configurations (see Figure 3), experiments were performed on a Leica SP8 microscope (as above) at the indicated emission filter settings. Acquisition was set at a scanning speed of 600 Hz and pinhole aperture was set at 3 AU.

### C. elegans - image processing and quantification

In some cases (Figure 1 (except G), Figure 2, Figure 4, Supplemental Figure S2 and S3), images of embryos/samples were taken alongside a local background image (with no samples in the field of view), which was subtracted from the image prior to analysis. This step can usually be omitted without much detriment, as an even and consistent background signal can be factored into the calibration parameters. However, background subtraction may improve images in cases where the background signal is uneven or variable. In some cases, a median filter (diameter 1 px) was applied before incorporation in figures.

Whole-embryo fluorescence intensities are defined as the mean pixel intensity within a manually defined region of interest (ROI) encompassing the embryo.

Individual cross-membrane profiles were extracted by taking 50-pixel line profiles perpendicular to and centered on the plasma membrane, using bicubic spline interpolation. Profiles were taken at pixel-width intervals around the circumference of the embryo, and averaged over the posterior-most (Figure 2E) or anterior-most (Figure 2F, 2G) 30% of the embryo’s circumference.

### SAIBR - Autofluorescence correction

#### 2 Channels

Fluorescence signal in the GFP Channel (G_Observed_) of an image can be described as a linear sum of true GFP (G_GFP_) and autofluorescence (G_AF_) contributions:

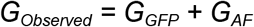

The aim of this procedure is to estimate G_AF_ in a given image, either on a whole-sample or pixel-by-pixel basis, so that this can be subtracted away and G_GFP_ determined. Our method exploits the fact that autofluorescence has a broad emission spectrum, and therefore can be captured in a red-shifted AF Channel (A):

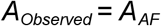

We assume the contribution of GFP to this channel to be very small (A_GFP_ ~ 0), owing to the narrow emission spectrum of GFP (however this assumption can be relaxed, as described in the section *GFP spillover correction*). Assuming that autofluorescence can be described as a single component with a characteristic emission spectrum, G_AF_ and A_AF_ (= A_Observed_) are expected to be linearly proportional:

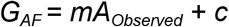

Therefore, provided that the appropriate inter-channel conversion factors (m and c) are known, G_AF_ can be calculated from A_Observed_ using the above equation. Parameters m and c can be determined by performing linear regression on data from GFP-free samples (for which G_Observed_ = G_AF_), as described in the section *Calculation of inter-channel correction factors*. The c parameter is included to account for potential differences in background intensity between the two channels.

#### 3 Channels

The above analysis can break down in samples containing red fluorophore, which can be weakly excited by 488 nm lasers, therefore adding an extra signal component to the A channel (A_RFP_):

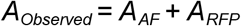

We can account for this contribution by using the RFP Channel (R), which is specific for RFP fluorophore (R_GFP_ ~ 0) and typically free of autofluorescence (R_AF_ ~ 0), as an independent readout of RFP levels:

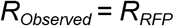

As A_RFP_ and R_RFP_ (= R_Observed_) are expected to be linear proportional, A_AF_, and therefore G_AF_ can be described as linear functions of A_Observed_ and R_Observed_:

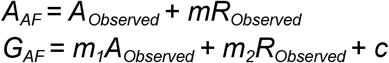

Therefore, provided that the appropriate inter-channel conversion factors (m_1_, m_2_ and c) are known, G_AF_ can be calculated from A_Observed_ and R_Observed_ using the above equation. The determination of these parameters is described below. As before, the c parameter is included to account for potential differences in background intensity between the channels.

#### Calculation of inter-channel correction factors

For two-channel SAIBR, correction parameters were calculated by performing the following linear regression on data from unlabeled samples (for which G_Observed_ = G_AF_):

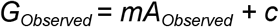

Where a red fluorophore is present, bleedthrough into the A channel must also be accounted for by using a three-channel method. These parameters were obtained by performing multiple linear regression using three-channel data from appropriate RFP-only samples:

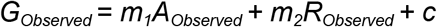

Note that in this study we used two methods to provide data for the regressions for *C. elegans* embryos: whole-embryo and pixel-by-pixel. In the whole embryo method G_Observed_ and A_Observed_ (and R_Observed_) represent mean whole-embryo intensity measurements for a series of embryos (see section *Quantification of whole-embryo fluorescence intensities*). In the pixel-by-pixel method G_Observed_ and A_Observed_ (and R_Observed_) represent pixel values taken from manually selected ROIs covering the entire embryo and a portion of the background, pooled from at least three embryos. We noted that using raw pixel values yielded relatively low correlations between channels, but that correlations increased by first applying a Guassian filter to the images to reduce salt and pepper noise (Supplemental Figure S2). A Gaussian radius of between 1 and 2 pixels was used for all analysis in this study. Linear regressions were performed using an ordinary least squares method. The plugin uses the latter pixel method.

Note that calibration samples were taken contemporaneously with test samples and imaged under identical conditions as changes to excitation / emission parameters will necessarily alter the parameters of the correction function.

#### GFP spillover correction

The long tail of the GFP emission spectrum means that a small fraction of GFP emission will appear in the AF Channel. This spillover of GFP signal into the AF Channel will artificially inflate A_AF_ and therefore result in oversubtraction upon application of SAIBR. However, because the magnitude of this effect is always proportional to GFP concentration, it simply rescales the magnitude of *G_GFP_* and thus can be safely ignored for normalized data or for relative comparisons between different GFP-containing samples. Alternatively, the magnitude of this effect can be measured and a correction applied. See Supplemental Figure S3 for additional details.

### Additional Methods - Starfish oocytes

Starfish (*P. miniata*, also known as *A. miniata*) oocyte collection and injection was performed as described before (Borrego-Pinto et al., 2016a; Terasaki, 1994). Briefly, starfish were obtained from Southern California Sea Urchin Co., Marinus Scientific, South Coast Bio-Marine, or Monterey Abalone Co. and maintained in seawater tanks at 16°C at the European Molecular Biology Laboratory (EMBL) Marine Facility. The mRNA encoding fluorescent mother centriolar Odf2-mEGFP (https://www.ncbi.nlm.nih.gov/nuccore/1040843242) (Borrego-Pinto et al., 2016b) was injected the day before imaging, while Cy3-tubulin (gift from the Nedelec laboratory, EMBL, Heidelberg, Germany) was injected shortly before imaging. After meiotic maturation with 10 μM 1-methyladenine (Acros Organics), oocytes were imaged on a Leica SP5 confocal microscope, as described in (Borrego-Pinto et al., 2016b). Sequential scanning was performed: in a first scan, 488 nm excitation was coupled to both mEGFP and red-shifted emission channels to record Odf2-mEGFP and autofluorescence, respectively. In a second scan, 561 nm excitation was combined with the same red-shifted emission channel to record Cy3-Tubulin fluorescence.

### Additional Methods - S. pombe

*Schizosaccharomyces pombe* (*S. pombe*) cells were grown in YES (yeast extract with supplements) medium overnight at 30°C. Prior to imaging, 1 mL *S. pombe* cell culture with OD_595nm_ 0.4-0.6 was concentrated to 50 μL after centrifugation at 1500x g, 30 sec. 2 μL of cell suspension were loaded under a 22 x 22 mm glass coverslip (VWR, thickness: 1.5). Spinning disk confocal images of *S. pombe* were captured with an Eclipse Ti-E inverted microscope fitted with Yokogawa CSU-X1 spinning disk confocal scanning unit, 600 series SS 488 nm, SS 561 nm lasers, single band filters FF01-525/50-25 and FF01-617/73-25 (Semrock Brightline), Nikon CFI Plan Apo Lambda 100x (NA = 1.45) oil objective and Andor iXon Ultra U3-888-BV monochrome EMCCD camera. Image acquisition was controlled by Andor IQ3 software.

## Contributions

Conceptualization: N.T.L.R., T.B., N.W.G.; Methodology: N.T.L.R., T.B.; Formal analysis: N.T.L.R., T.B.; Investigation: N.T.L.R., T.B., J.B.-P., K.N., Y.G., S.F.; Resources: N.T.L.R., T.B., N.H.; Writing - original draft preparation: N.T.L.R., T.B., N.W.G.; Writing - review and editing: N.T.L.R., T.B., N.W.G.; Supervision: N.W.G.; Project administration: N.W.G.; Funding acquisition: N.W.G.

## Acknowledgements

We thank Josana Rodriguez, Ruben Schmidt, and Donald Bell for comments on the manuscript and/or pilot testing of the SAIBR plugin. Strains and/or reagents were graciously provided by Dan Dickinson and Ken Kemphues. Additional strains were provided by the Caenorhabditis Genome Center (CGC), which is funded by NIH Office of Research Infrastructure Programs (P40 OD010440). Confocal imaging was performed with assistance of the Crick Advanced Light Microscopy (CALM) STP. Some experiments were performed in the labs of Peter Lenart (EMBL, MPI-BPC) and Snezhka Oliferenko (Francis Crick Institute). This work was supported by the Francis Crick Institute, which receives its core funding from Cancer Research UK (FC001086), the UK Medical Research Council (FC001086), and the Wellcome Trust (FC001086).

This research was funded in whole, or in part, by the Wellcome Trust (FC001086). For the purpose of Open Access, the author has applied a CC BY public copyright licence to any Author Accepted Manuscript version arising from this submission.

## Competing Interests

No competing interests declared.

## Data Availability

Source code and documentation for the plugin are available at https://github.com/goehringlab/saibr_fiji_plugin under a CC BY 4.0 license. Original image datasets available upon reasonable request from the corresponding author.

---

**Box 1.**
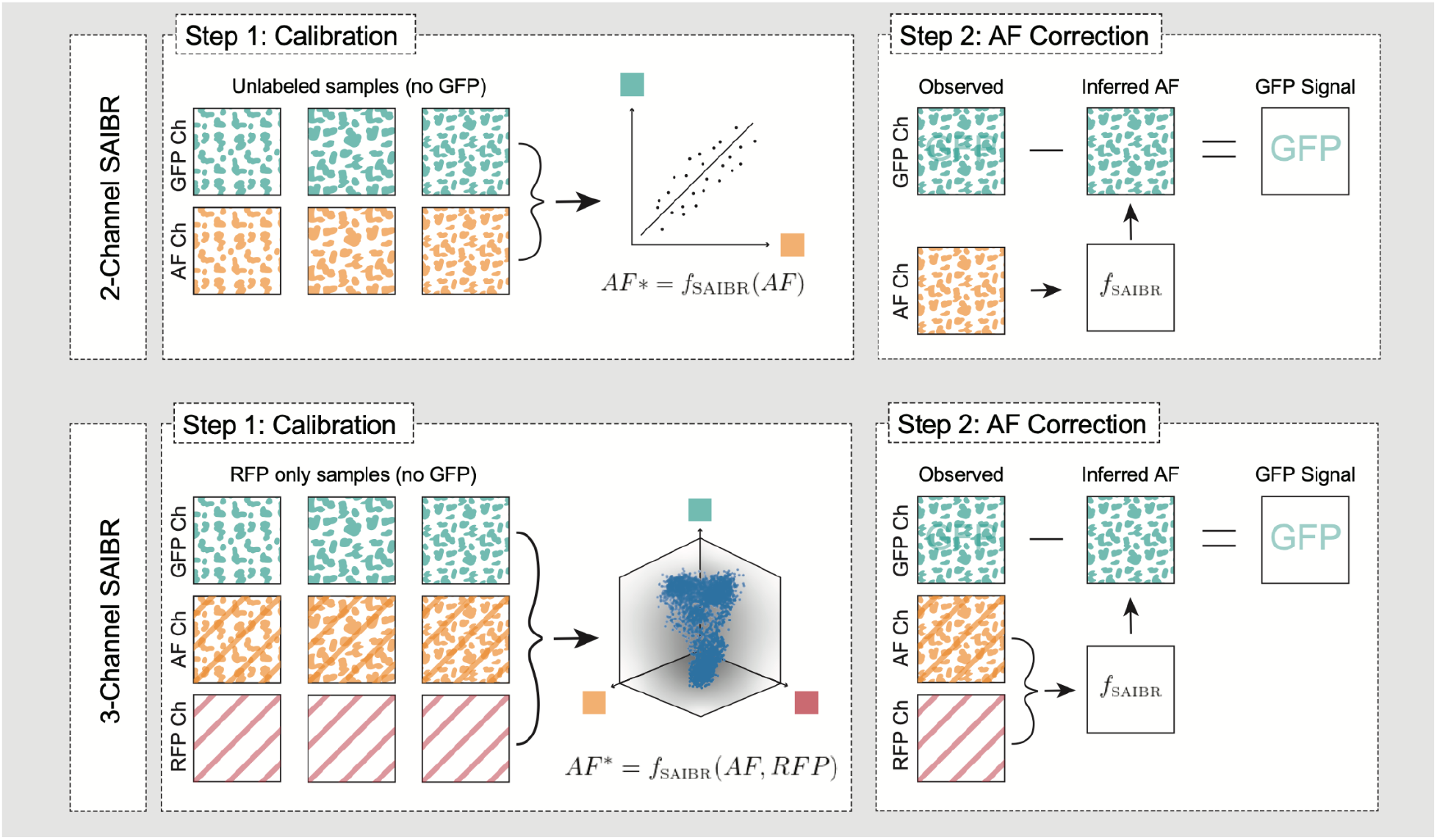
Schematic summary of SAIBR workflow. The first step for 2-Channel SAIBR is to define the correction function *f_SAIBR_*. *f_SAIBR_* is defined by imaging multiple unlabeled calibration samples in which all fluorescence signal arises from autofluorescence and performing a linear regression on fluorescence values in the GFP and AF Channels. In step 2, to isolate GFP signal in GFP-labeled samples, measured AF Channel signal is used to infer AF signal in the GFP Channel using *f_SAIBR_* and inferred AF signal is then subtracted from the observed GFP Channel signal to yield the “true GFP” signal. In 3-Channel SAIBR, one must compensate for red fluorophore (e.g RFP) signal bleedthrough into the AF Channel. Therefore, for 3-Channel SAIBR, *f_SAIBR_* is defined by imaging samples expressing only RFP and performing a multiple linear regression to correlate observed AF and RFP Channel signals to signal in the GFP Channel. One can then use *f_SAIBR_* to infer AF in GFP-labeled samples based on AF and RFP Channel signal and subtract from observed GFP Channel signal to obtain the true GFP signal.

## Notes

https://github.com/goehringlab/saibr_fiji_plugin

### Competing Interest Statement

The authors have declared no competing interest.

