## Supplemental Table S1 for "*SAIBR*: A simple, platform-independent method for spectral autofluorescence correction"

### Table S1. Key reagents and resources

#### Oligonucleotides

| Description | Sequence |
|---|---|
| *lgl-1* crRNA #1 | CTCCGAACTACAGCGAAGTA |
| *lgl-1* crRNA #2 | ACGGTGAATTTGAACTTTCG |
| GFP fwd ODN | ATGAGTAAAGGAGAAGAATTGTT |
| GFP rev ODN | CTTGTAGAGCTCGTCCATTCC |
| *lgl-1* HDR fwd ODN | TCTTCATTTGAGTCGAGGAAGTTTATAGACCCAATCGACGTAATGGCACTAAACTCGAGTTGTTTATCACGTGAT GGTGACGTGGCATTCCTCATCAAATCCTCCGAACTACAGCGAAGTACAGTAAACCTCAATTTCAGAGGTTGGAG CAGTACGCACAAGTCAATGAGTAAAGGAGAAGAATTGTT |
| *lgl-1* HDR rev ODN | CAAAAAAGGCAAAGACCGAGGGCAAATAAATAACATAATAAAGTTTAAAAAAAACCACCATTTCAAACAAAATTAA TATATATCAACAGGAAAACGATTTTTAAAAAAAATGCATCTACTTGTAGAGCTCGTCCATTCCG |

#### *C. elegans* strains

| Strain | Description | Genotype | Source | Method |
|---|---|---|---|---|
| N2 | Wild type | Wild type | CGC | Wild type |
| KK1248 | PAR-6::GFP | *par-6(it310[par-6::gfp]) I* | CGC/Ken Kemphues | CRISPR/Cas9 |
| KK1216 | PAR-3::GFP | *par-3(it298[par-3::gfp]) III* | CGC/Ken Kemphues | CRISPR/Cas9 |
| NWG0285 | LGL-1::GFP | *lgl-1(crk66[lgl-1::gfp]) X* | This paper | CRISPR/Cas9 |
| DG4190 | GFP::cdc25.3 | *cdc-25.3(tn1712[gfp::3xflag::cdc-25.3]) III* | CGC/David Greenstein | CRISPR/Cas9 |
| LP216 | PAR-6::mNG | *par-6(cp45[par-6::mneongreen::3xFlag + LoxP unc-119(+) LoxP]) I; unc-119(ed3) III* | CGC/Dan Dickinson | CRISPR/Cas9 |
| TH209 | mCh::PAR-2 | *unc-119(ed3) III; ddIs31[pie-1p::mcherry::par-2; unc-119(+)]* | Brangwynne et al., 2009 | Biolistic transformation |
| NWG0033 | mCh::MEX-5 | *unc-119(ed3) III; axIs1731[pie-1p::mcherry::mex-5::pie-1 3'UTR + unc-119 (+)]* | Derived from JH2840 (CGC/Geraldine Seydoux) | Biolistic transformation |
| NWG0119 | PAR-6::GFP, mCh::MEX-5 | *par-6(it310[par-6::gfp]) I; unc-119(ed3) III; axIs1731[pie-1p::mcherry::mex-5::pie-1 3'UTR + unc-119 (+)]* | Parent strains: KK1248, NWG0033 | CRISPR/Cas9 and biolistic transformation |
| BOX241 | PAR-6::mCh | *par-6(mib25[par-6::mcherry-LoxP]) I* | Mike Boxem | CRISPR/Cas9 |
| NWG0290 | PAR-6::mCh, LGL-1::GFP | *par-6(mib25[par-6::mcherry-LoxP]) I; lgl-1(crk67[lgl-1::gfp]) X* | Parent strains: BOX241, NWG0286 (this paper) | CRISPR/Cas9 |
| KK1262 | PAR-1::GFP | *par-1(it324[par-1::gfp::par-1 exon 11a]) V* | CGC/Ken Kemphues | CRISPR/Cas9 |
| LP306 | GFP::PH | *cpIs53 [mex-5p::gfp-C1::plc(delta)-PH::tbb-2 3'UTR + unc-119 (+)] II; unc-119(ed3) III* | CGC/Bob Goldstein | Insertion at ttTi5605 MosI site using CRISPR/Cas9 |

#### *S. pombe* strains

| Strain | Description | Genotype | Source | Method |
|---|---|---|---|---|
| SO2865 | Untagged *S. pombe* | ade6-210 ura4-D18 leu1-32 h+ | Snezhana Oliferenko | Endogenous fluorescence tagging |
| SO8304 | Nem1::mNG | nem1-mNeonGreen::kanR ade6-216 ura4-D18 leu1-32 h- | Snezhana Oliferenko | Endogenous fluorescence tagging |
