## Supplemental Material and Figures S1-S3 for "*SAIBR*: A simple, platform-independent method for spectral autofluorescence correction"

Supplemental Figure 1 - Rodrigues, Bland, et al. (column-wide figure)

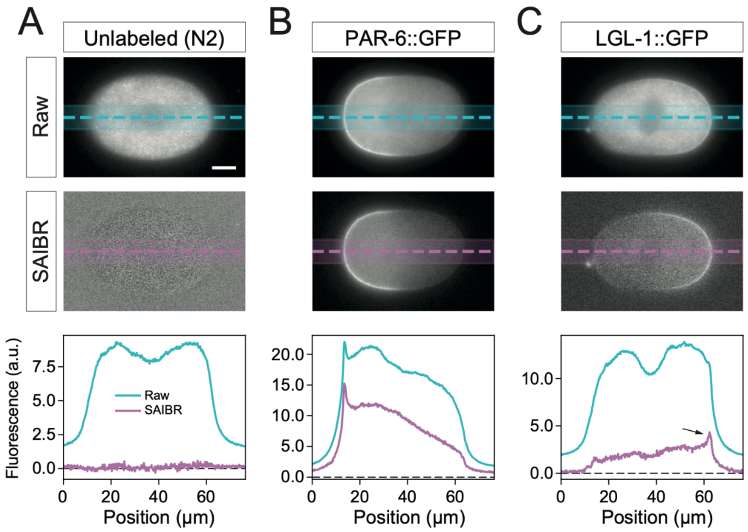


**Supplemental Figure S1. Spectral AF correction using widefield fluorescence microscopy for LGL-1::GFP.** Raw (top) and SAIBR-corrected (middle) midplane images of **(A)** unlabeled wild-type (N2) control, **(B)** PAR-6::GFP (strain: KK1248), and **(C)** LGL-1::GFP (strain: NWG0285) zygotes imaged with widefield fluorescence microscopy. (bottom) Quantification of fluorescence linescans taken across the embryos as indicated. Arrow highlights plasma membrane signal in LGL-1::GFP linescan that is practically undetectable in uncorrected images. Scale Bars = 10 µm.

Supplemental Figure 2 - Rodrigues, Bland, et al. (page-wide figure)

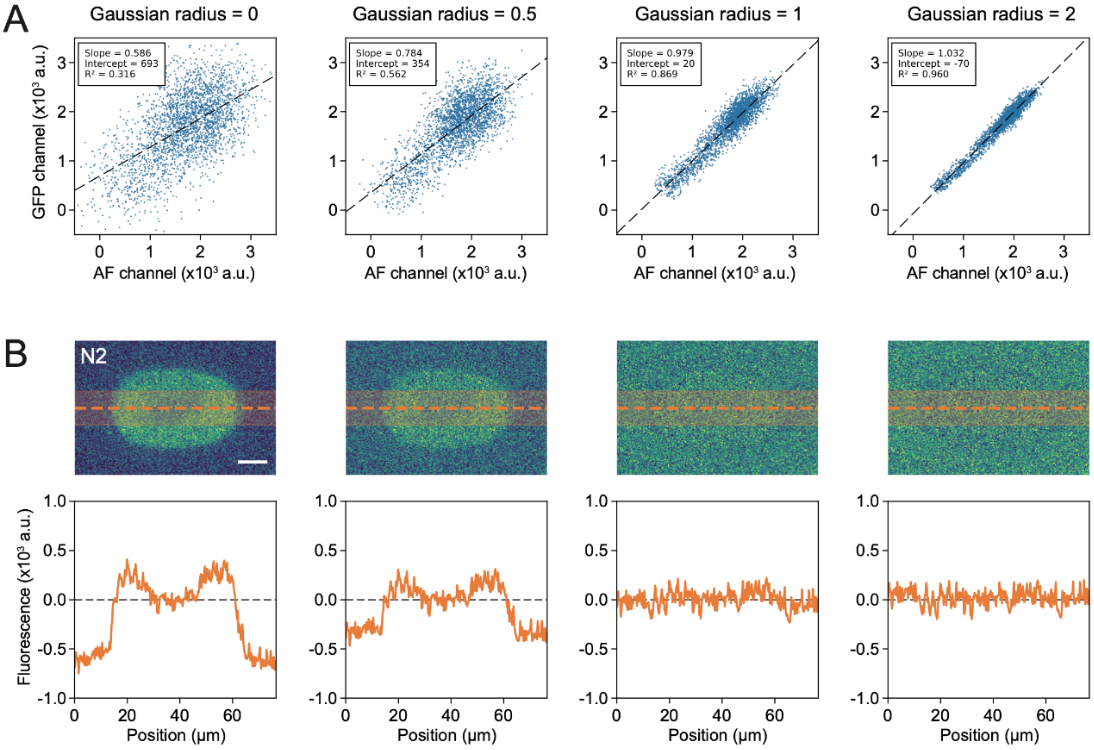


**Supplemental Figure S2. Suppression of shot noise improves per-pixel AF to GFP Channel correlation. (A)** Per-pixel plots of AF vs GFP Channel signal for images of unlabeled embryos subject to gaussian blur of indicated radius. Linear fit shown as dashed line with parameters in inset indicating improved fit with increasing gaussian radius. **(B)** Results on AF correction of unlabeled N2 embryos using the linear fits in (A). Processed images (top) and line scan across the central region of the embryo (bottom) show improved suppression of AF with increasing Gaussian radius.

Supplemental Figure 3 - Rodrigues, Bland, et al. (page-wide figure)

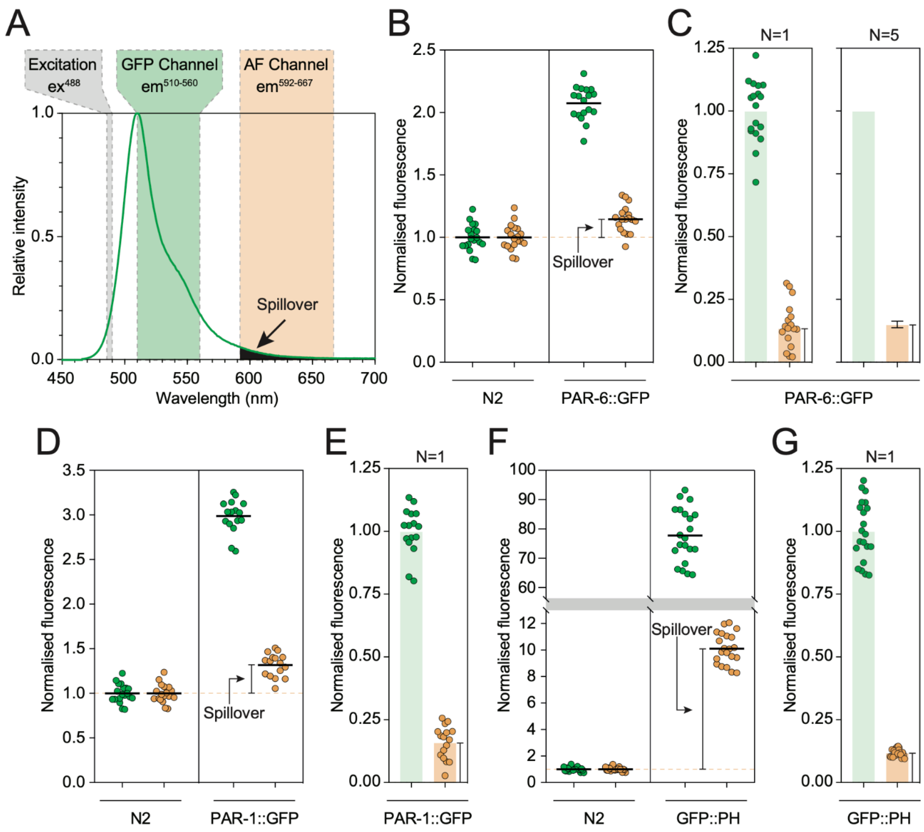


**Supplemental Figure S3. Quantification of GFP spillover into AF Channel. (A)** The long tail of the GFP emission spectrum (data from searchlight.semrock.com) means that some GFP signal will appear in the AF Channel ($em^{592\text{-}667}$). Predicted spillover in the AF Channel shaded in black. **(B-G)** Quantification of the magnitude of GFP spillover into the AF Channel reveals it is a consistent fraction of GFP signal (here ~15%). Spillover was detected by comparing AF Channel signal from unlabeled and GFP-expressing embryos (GFP Channel - green, AF Channel - orange). Mean fluorescence signal for the indicated strains/channels (B, D, E) and spillover as a fraction of normalized GFP signal (C, E, G) shown. Note for PAR-6::GFP (strain: KK1248), we replicated this measure five times and show both a representative replicate and mean±SD for all five replicates. A single replicate was performed each for PAR-1::GFP (strain: KK1262) and GFP::PH (strain: LP306) for comparison. While this spillover of GFP signal into the AF Channel will result in oversubtraction when applying SAIBR, because the magnitude of this effect is always proportional to GFP concentration, it simply rescales the magnitude of the obtained GFP value and thus is irrelevant for normalized data or for comparisons between different GFP-containing samples.

**Supplemental Table S1.** Reagents and Resources

**Supplemental Movie 1.** Timelapse of *C. elegans* embryo expressing LGL-1::GFP, PAR-6::mCherry at the endogenous loci from the 1- to 4-cell stage highlighting differences between uncorrected and corrected images for the GFP channel. DIC, mCherry, raw GFP and SAIBR images shown. Frame rate 1/min.
